# FoldToken3: Fold Structures Worth 256 Words or Less

**DOI:** 10.1101/2024.07.08.602548

**Authors:** Zhangyang Gao, Cheng Tan, Stan Z. Li

## Abstract

Protein structure tokenization has attracted increasing attention in both protein representation learning and generation. While recent work, like FoldToken2 and ESM3, has achieved good reconstruction performance, the compressoin ratio is still limited. In this work, we propose FoldToken3, a novel protein structure tokenization method that can compress protein structures into 256 tokens or less and ensure the reconstruction quality comparable to FoldToken2. To the best of our knowledge, FoldToken3 is the most efficient, light-weight, and compression-friendly protein structure tokenization method. And it will benifit a wide range of protein structure-related tasks, such as protein structure alignment, generation, and representation learning. The work is still in progress and the code will be available upon acceptance.

## 1 Introduction

> “SE-(3) structure should not be special and difficult. Let’s lower the barrier.”
>
> – Our Goal

The SE-(3) nature of protein structure has posed long-term challenges in representation learning and generation, requiring researchers to carefully design the invariant encoder [5, 9] and equivariant decoder [1, 14]. Theses models are usually computationally expensive and difficult to understand for non-AI experts. Inspired by the success in computer vision [4] and multimodal learning [17], tokenizing equivariant structures as invariant discrete tokens has emerged as a promising direction. This approach simplifies model design by leveraging current NLP and CV models and enhances multimodal capabilities through the use of a unified fold language.

Existing tokenization methods are limited in compression ratio. For example, FoldToken2 [8] and ESM3 [10] have the codebook size of 65536 and 4096, respectively. The large codebook size pose challenges in several aspects: (1) hard to analyze all the structure patterns; (2) leads to difficulty in downstream generative tasks, as the predictive space is large and similar code vectors could confuse each other. An open problem is: *how to compress the code space into less tokens while maintaining the reconstruction quality remains?*

FoldToken2 suffers from the lack of diverse code vectors due to unstable gradient. Regarding the quantifier, when the temperature parameter is extremly small, the unstable gradient causes codebook collapse, making code vectors to be highly similar. These similar code vectors can easily confuse each other, resulting in additional difficulty in generative tasks. Moreover, even small noise can drastically change the coding sequence, leading to inconsistency between similar proteins. Regarding the encoder, the unstable gradient will also disrupt the its representation capability, making the learned embeddings to be less informative.

FoldToken3 re-designs the vector quantization module to address above issues. Firstly, we propose a ‘partial gradient’ trick to allow the encoder and quantifier receive stable gradient no matter how the temperature is small. Secondly, we replace the ‘argmax’ operation as sampling from a categorical distribution, making the code selection process to be stochastic. The first innovation makes the model to be training stable and overcome the codebook collapse issue. The second innovation makes the codebook to be more diverse and robust to noise.

With the highly-optimized vector quantization module, we find that even a small codebook size can achieve comparable reconstruction quality to FoldToken2. Compared to ESM3, whose encoder and decoder have 30.1M and 618.6M parameters with 4096 code space, FoldToken3 has 4.31M and 4.92M parameters with 256 code space. In addition, we have reduced the code space to 256 or less, under 0.4% of the FoldToken2 code space. We believe that FoldToken3 will benefit a wide range of protein structure-related tasks.

## 2 Method

### 2.1 Overall Framework

As shown in Fig. 1, the overall framework keeps the same as FoldToken1 [6, 8] and FoldToken2 [7], including encoder, quantifier and decoder. From FoldToken2 to FoldToken3, we make the following improvements:

**Figure 1:**
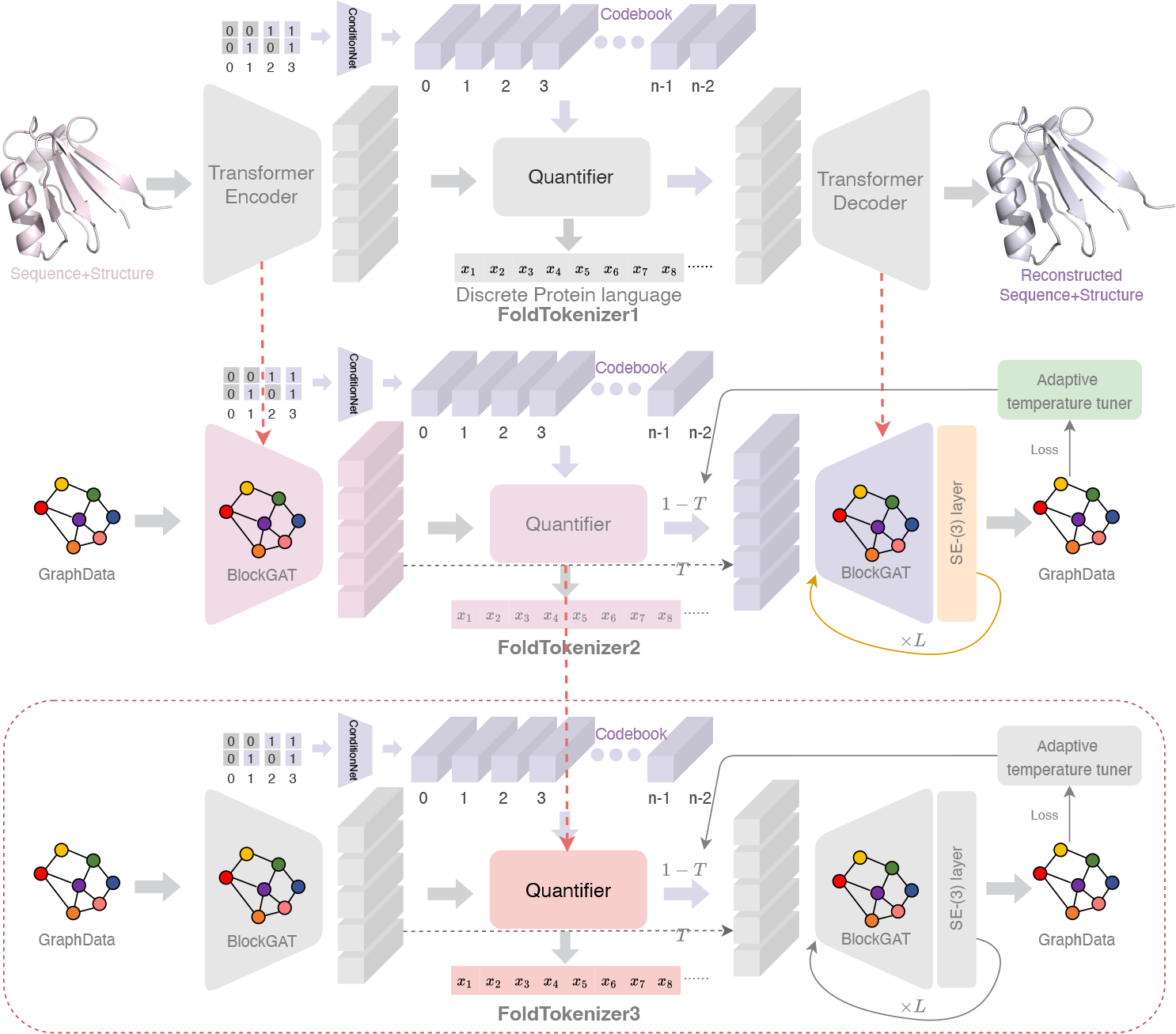
The overall framework of FoldTokenizer3, which contains contains encoder, quantifier, and decoder. We use BlockGAT to encoder protein structures as invariant embeddings, SoftCVQ to quantize the embeddings into discrete tokens, and SE-(3) layer to recover the protein structures iteratively.

#### 1. Stochastic Selection

We replace the “argmax” operation with sampling from a categorical distribution to select the nearest code vector.

#### 2. Reparameterization

We reparameterize the categorical random variable to allow the quantifier to allow the network to optimise the distribution parameters.

#### 3. Stablize Gradient

We introduce the trick of ‘partial gradient’ to stablize the gradient of the quantifier and encoder.

### 2.2 Background of Vector Quantization

#### Vector Quantization Problem

The quantifier *Q* : ***h***↦ *z* converts continuous embedding ***h*** as a discrete latent code *z*, named VQ-ID. The de-quantifier 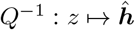 recovers continuous embedding 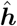 from *z*. The problem can be formulated as

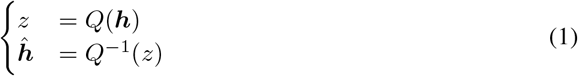

Readers should read FoldToken1 [8] to better understand the limitations of vanilla vector quantifier (VVQ) and lookup-free quantifier (LFQ):

#### Limitations of VVQ

The vanilla quantifier directly copies the gradient of 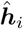 to ***h***_*i*_, i.e., 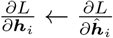 resulting in the **gradient mismatching** between embedding 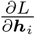 and ***h***_*i*_. In addition, the higher reconstruction quality does not necessarily lead to better generation performance on downstream tasks. For example, [16] points out that reconstruction and generation may be contradicted: *Enlarging the vocabulary can improve reconstruction quality. However, such improvement only extends to generation when the vocabulary size is small, and a very large vocabulary can actually hurt the performance of the generative model*. Why does the contradiction occur? We attribute the intrinsic reason to the **large class space**. We summarize the limitations as follows:

##### 1. Gradient Mismatching

As the VQ-ID 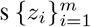 are non-differentiable, the vanilla quantifier directly copies the gradient of 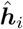 to ***h***_*i*_, i.e., 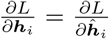. However, 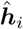 generally does not equal ***h***_*i*_, resulting in the mismatching between embedding ***h***_*i*_ and its gradient 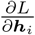.

##### 2. Large Irrelevant Class Space

Regarding the generative model, each VQ-ID *z*_*i*_ represents a class index. The discrete and continuous representations exhibit no association between (***h***_*i*_-***h***_*j*_) and (*z*_*i*_-*z*_*j*_), indicating that the generative model should accurately predict the exact *z*_*i*_, despite the high similarity between *z*_*i*_ and *z*_*j*_. As the codebook size could be large, predicting VQ-ID without considering their associations poses generation challenge.

#### Limitations of LFQ

LFQ fails to address the issue of gradient mismatching and and introduces a new challenge of information bottleneck. Typically, the codebook space is chosen from options such as 2^8^ (int8), 2^16^ (int16), 2^32^(int32), and 2^64^(int64) due to accommodate the required bits for storing a VQ-ID. However, for a fair comparison to VVQ and effective data compression, only 2^8^ and 2^16^ are considered, resulting in a hidden space size of 8 or 16 in LFQ. The low dimensionality poses the problem of information bottleneck in the enc-decoder model, which hampers accurate reconstruction.

### 2.3 Binary Stochastic Quantifier (Novel Part)

Unlike FoldToken1 [6, 8] and FoldToken2 [7] that use SoftCVQ, we introduce a novel quantifier, called Binary Stochastic Quantifier (BSQ), to quantize the embeddings.

Given the decimal integer *z*_*i*_ and the codebook size *m*, we represent *z*_*i*_ in binary form ***b***_*i*_ with length log_2_(*m*). For example, if *m* = 4, we have ***b***_0_ = [0, 0], ***b***_1_ = [0, 1], ***b***_2_ = [1, 0], ***b***_3_ = [1, 1]. Instead of using `nn.Embedding` layer to encode *z*_0_ and *z*_1_ independently, we use a MLP, called `ConditionNet`, to project ***b***_0_ and ***b***_1_ to code vectors ***v***_0_ and ***v***_1_ to consider their inherent correlations in each bit position. The binary code ensure the propsoed model share the advantage of LFQ in terms of robust generation and relevant semantics; the `ConditionNet` enlarges the dimensionlity of the binary code to overcome the LFQ’s shortcoming of information bottleneck.

Formally, we explain the binary code and `ConditionNet` as

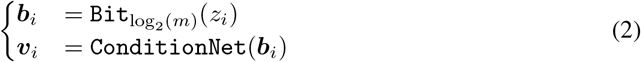

where ***v***_*j*_ is the *j*-th code embedding. The ConditionNet : 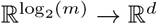 is a MLP. If the codebook size is 2^10^, the MLP projects 1024 16-dimension binary vectors into 1024 *d*-dimension code vectors.

The key problem is: *how to replace latent embedding* ***h***_*i*_ *with the most similar token embedding* ***v***_*j*_ *in a differential way*.

#### Find Neighbor

In vanilla vector quantization, they use the nearest neighbor algorithm to find the most similar code vector, which is non-differential. In this paper, we take the selection process as sampling from a multi-class distribution *z*_*i*_ ∼ `Mult`(***p***_*i*_):

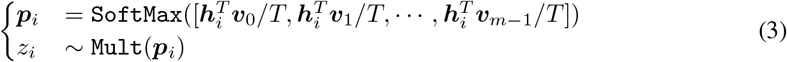

and then, we optimize ***p***_*i*_ in a differential way. The temperature parameter *T* controls the softness of the attention query operation. When *T* is large, the attention weights will be close to uniform; otherwise, the attention weights will be close to one-hot.

#### Optimize Neighbor

In Eq. 3, the sampling operation is non-differentiable, and we use a reparameterization trick to optimize ***p***_*i*_:

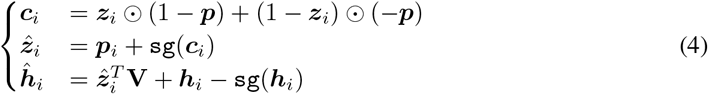

where `sg`(·) is the stop gradient operation, ⊙ is the element-wise multiplication, and ***z***_*i*_ ∈ ℝ^*m*^ is the onehot version of *z*_*i*_. The first two equations reparameterize ***z***_*i*_ as 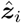 to allow ***p***_*i*_ get gradient, inspired by [11]; the third equation allows ***h***_*i*_ to get direct gradient for optimizing the encoder. 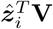 operation selects the *z*_*i*_-th code vector using the onehot 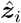.

#### Teacher Guidance

To accelerate training convergence, we randomly copy encoder output ***h***_*i*_ as decoder input 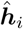 in probability *T* :

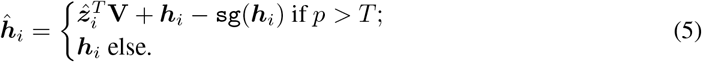

where *p*∼ *U* (0, 1) is sampled from uniform distribution. When *T* = 1.0, the vector quantization module is skipped, allowing the encoder-decoder to be easily optimized. When *T* = 0.0, the vector quantization module is fully used. For values of 0.0 *< T <* 1.0, the shortcut feature guides the vector quantization model to learn code vectors that align with the encoder inputs. The adaptive temperature scheduler is:

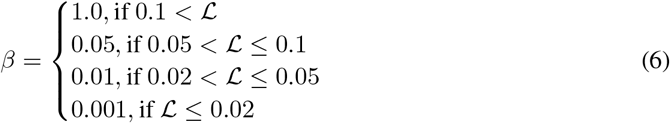

#### Stable Gradient

The scaled softmax operation in Eq. 4 bridges the continual model (*T >* 0) to discrete vector quantization (*T* = 0); thus allowing precise gradient computation rather than gradient mismatch in the VVQ. During training, we gradually anneal the temperature from 1.0 to 1e-8; however, the gradient of the scaled softmax tend to explode when *T* is small:

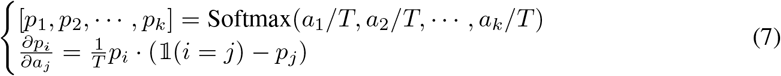

The unstable gradient would lead to representation and codebook collapses, as the `ConditionNet` and encoder parameters collapse after one step of updating a large gradient. To overcome the issue, we introduce the trick of ‘partial gradient’:

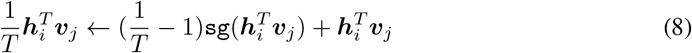

where the first term is stoped gradient and only the second term contribute to gradient computation. Obviously, Eq.8 has the same forward behavior like Eq.4 while the gradient is stable and do not affected by the extreme small value of *T*.

### 2.4 Invariant Graph Encoder (Same as FoldToken2)

Due to the rotation and translation equivariant nature, the same protein may have different coordinate records, posing a challenge in learning compact invariant representations for the same protein. Previous works [5, 13, 3, 9] have shown that the invariant featurizer can encode the invariant structure patterns, and we follow the same road: representing the protein structures as a graph consisting of invariant node and edge features. Then we use the BlockGAT [9] to learn high-level representations.

#### Frame-based Block Graph

Given a protein 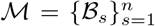 containing *n* blocks, where each block represents an amino acid, we build the block graph 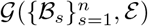 using kNN algorithm. In the block graph, the *s*-th node is represented as ℬ_*s*_ = (*T*_*s*_, ***f***_*s*_), and the edge between (*s, t*) is represented as ℬ_*st*_ = (*T*_*st*_, ***f***_*st*_). *T*_*s*_ and 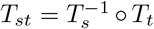 are the local frames of the *s*-th and the relative transform between the *s*-th and *t*-th blocks, respectively. ***f***_*s*_ and ***f***_*st*_ are the node and edge features.

#### BlockGAT Encoder

We use the BlockGAT [9] layer *f*_*θ*_ to learn block-level representations:

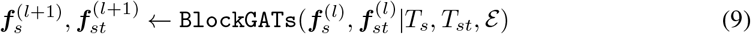

where 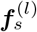 and 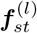 represent the input node and edge features of the *l*-th layer. *T* _*s*_ = (*R* _*s*_, ***t*** _*s*_) is the local frame of the *s*-th block, and 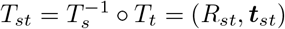 is the relative transform between the *s*-th and *t*-th blocks. *T*_*s*_, *T*_*st*_, 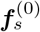 and 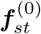are initialized from the ground truth structures using the invariant featurizer proposed in UniIF [9].

### 2.5 Equivariant Graph Decoder (Same as FoldToken2)

Generating the protein structures conditioned on invariant representations poses significant challenges in computing efficiency. For example, training well-known AlphaFold2 from scratch takes 128 TPUv3 cores for 11 days [15]; OpenFold takes 50000 GPU hours for training [2]. In this work, we propose an efficiency plug-and-play SE(3)-layer that could be added to any GNN layer for structure prediction. Thanks to the simplified module of the SE(3)-layerand BlockGAT with sparse graph attention, we can train the model on the entire PDB dataset in 1 day using 8 NVIDIA-A100s.

#### SE-(3) Frame Passing Layer

We introduce frame-level message passing, which updates the local frame of the *s*-th block by aggregating the relative rotation *R*_*s*_ and translation ***t***_*s*_ from its neighbors:

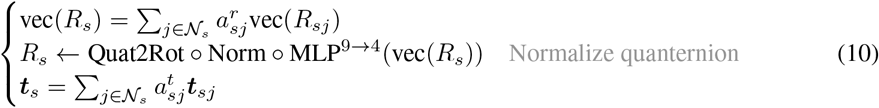

where 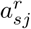 and 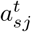 are the rotation and translation weights, and 𝒩_*s*_ is the neighbors of the *s*-th block. vec(·) flattens 3 × 3 matrix to 9-dimensional vector. MLP^9*→*4^(·) maps the 9-dim rotation matrix to 4-dim quaternion, and Norm(·) normalize the quaternion to ensure it represents a valid rotation. Quat2Rot(·) is the quaternion to rotation function. We further introduce the details as follows:

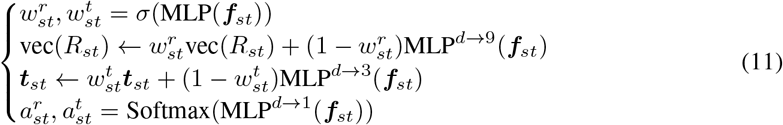

where 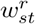 and 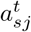 are the updating weights for rotation and translation, 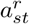 and 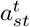 are the attention weights. The propose SE-(3) layer could be add to any graph neural network for local frame updating.

#### Iterative Refinement

We propose a new module named SE-(3) BlockGAT by adding a SE-(3) layer to BlockGAT. We stack multi-layer SE-(3) BlockGAT to iteratively refine the structures:

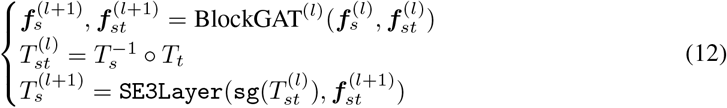

where `sg`(·) is the stop-gradient operation, and `SE3Layer`(·) is the SE-(3) layer described in Eq.11. Given the predicted local frame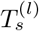, we can obtain the 3D coordinates by:

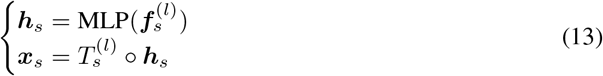

### 2.6 Reconstruction Loss (Same as FoldToken2)

Inspired by Chroma [12], we use multiple loss functions to train the model. The overall loss is:

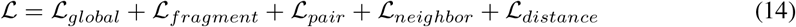

To illustrate the loss terms, we define the aligned RMSD loss as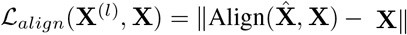, given the the ground truth 3D coordinates **X** ∈ ℝ^*n*,3^ and the predicted 3D coordinates 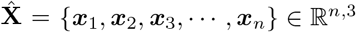. The global, fragment and pair loss are defined by the aligned MSE loss, but with different input data shape:

- **Global Loss**: **X** with shape [*n*, 4, 3]. RMSD of the global structure.
- **Fragment Loss**: **X** with shape [*n, c*, 4, 3]. RMSD of *c* neighbors for each residue.
- **Pair Loss**: **X** with shape [*n, K, c* · 2, 4, 3]. RMSD of *c* neighbors for each kNN pair.
- **Neighbor Loss**: **X** with shape [*n, K*, 4, 3]. RMSD of *K* neighbors for each residue.

where *n* is the number of residues, *c* = 7 is the number of fragments, *K* = 30 is the number of kNN, 4 means we consider four backbone atoms {*N, CA, C, O* }, and 3 means the 3D coordinates. The distance loss is defined as the MSE loss between the predicted and ground truth pairwise distances:

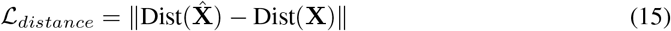

where Dist(**X**) ∈ ℝ^*n,n*^ is the pairwise distance matrix of the 3D coordinates **X**. We apply the loss on each decoder layer, and the final loss is the average, whcih is crucial for good performance.

## 3 Experiments

We conduct systematic experiments to inspire further improvement in FoldToken2.

- **Single-Chain Benchmark (Q1)**: How well FoldToken3 perform on single-chain data?
- **Multi-Chain Benchmark (Q2)**: How well FoldToken3 perform on multi-chain data?
- **VQ Insights (Q3):** What can we learn from FoldToken3’s improvement?

### Multi-chain PDB Data for Training

We train the model using all proteins collected from the PDB dataset as of 1 March 2024. After filtering residues with missing coordinates and proteins less than 30 residues, we obtain 162K proteins for training. We random crop long proteins to ensure that the maximum length is 500. Protein complexes are supported by adding chain encoding features *c*_*ij*_ to the edge *e*_*ij*_: *c*_*ij*_ = 0, if *i* and *j* are in different chains; else *c*_*ij*_ = 1. We train FoldToken2 and FoldToken3 on the protein structure reconstruction task using the PDB dataset. The model is trained for up to 25 epochs with a batch size of 8, a learning rate of 0.001, and a padding length of 500.

#### 3.1 Single-Chain Benchmark (Q1)

##### Metrics

Regarding reconstruction, we evaluate the model using the average TMscore and aligned RMSD. In FoldToken2, we uses Kabsch algorithm to align the predicted structure to the ground truth structure; however, the aligned RMSD seems to be different to that of PyMol. We do not know what is the reason for this discrepancy. Finally, we use PyMol’s API for computing RMSD. We also introduce a similarity metric to evaluate the codebook diversity:

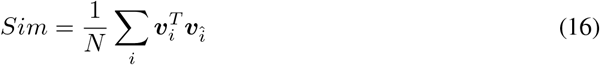

where ***v***_*i*_ and 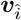 are the *i*-th and nearest neighbor code vectors, respectively. The similarity metric ranges from -1 to 1, with 1 indicating that there is always a very similar code vector for each one. To make discrete tokens distinguishable, the smaller the similarity, the better the diversity, and the easier it is to predict. Considering the extreme case where the similarity is 1, one code id can be replaced by that of its nearest neighbor without affecting the reconstruction quality, leading to inconsistent language representation. Also, high similarity indicates that the model do not robust to noise, as similar code vectors may be easily confused by each other.

##### Single-Chain Data for Evaluation

Following FoldToken2, we evaluate models on T493 and T116 datasets for head-to-head comparison, which contains 493 and 116 proteins, respectively. We also evaluate methods on 128 novel proteins released after the publication of AlphaFold3, called N128.

In Table. 1, we show the reconstruction results of FoldToken2 and conclude that:

**Table 1:** Single-chain Reconstruction Benchmark. FT1, FT2, and FT3 indicates FoldToken1 [6, 8], FoldToken2 [7], and FoldToken3, respectively. We also report the reconstruction results of ESM3 [10] for comprehensive understanding. When KNN is ‘full’, the approach uses full attention to consider all pairwise interactions.

| Model | Data | Config | | | | | TMScore ↑ | | | RMSD ↓ | | |
| --- | --- | --- | --- | --- | --- | --- | --- | --- | --- | --- | --- | --- |
|  |  | #Code | #Enc | #Dec | #Hid | #KNN | Avg | Max | Min | Avg | Max | Min |
| FT1 | T116 | 65536 | 12 | 12 | 480 | full | 0.77 | 0.96 | 0.39 | 3.31 | 24.53 | 0.52 |
| FT2 | T116 | 65536 | 8 | 8 | 128 | 30 | 0.97 | 0.99 | 0.90 | 0.52 | 0.76 | 0.30 |
| ESM3 | T116 | 4096 | 2 | 30 | 1024 | full | 0.97 | 0.99 | 0.76 | 1.97 | 13.27 | 0.01 |
| FT3 | T116 | 4096 | 8 | 8 | 128 | 30 | 0.95 | 0.98 | 0.88 | 0.64 | 0.98 | 0.30 |
| FT3 | T116 | 1024 | 8 | 8 | 128 | 30 | 0.93 | 0.97 | 0.83 | 0.73 | 1.20 | 0.26 |
| FT3 | T116 | 256 | 8 | 8 | 128 | 30 | 0.93 | 0.98 | 0.86 | 0.76 | 1.10 | 0.44 |
| FT3 | T116 | 128 | 8 | 8 | 128 | 30 | 0.91 | 0.96 | 0.82 | 0.87 | 1.36 | 0.46 |
| FT3 | T116 | 64 | 8 | 8 | 128 | 30 | 0.89 | 0.96 | 0.74 | 1.02 | 1.80 | 0.32 |
| FT1 | T493 | 65536 | 12 | 12 | 480 | full | 0.74 | 0.96 | 0.44 | 3.09 | 18.09 | 0.48 |
| FT2 | T493 | 65536 | 8 | 8 | 128 | 30 | 0.95 | 0.99 | 0.78 | 0.64 | 1.99 | 0.36 |
| ESM3 | T493 | 4096 | 2 | 30 | 1024 | full | 0.95 | 0.99 | 0.32 | 2.40 | 15.97 | 0.01 |
| FT3 | T493 | 4096 | 8 | 8 | 128 | 30 | 0.92 | 0.99 | 0.57 | 0.86 | 6.48 | 0.35 |
| FT3 | T493 | 1024 | 8 | 8 | 128 | 30 | 0.90 | 0.98 | 0.61 | 0.98 | 7.52 | 0.38 |
| FT3 | T493 | 256 | 8 | 8 | 128 | 30 | 0.90 | 0.98 | 0.52 | 1.03 | 6.14 | 0.38 |
| FT3 | T493 | 128 | 8 | 8 | 128 | 30 | 0.88 | 0.97 | 0.49 | 1.15 | 7.70 | 0.39 |
| FT3 | T493 | 64 | 8 | 8 | 128 | 30 | 0.85 | 0.96 | 0.45 | 1.33 | 7.47 | 0.47 |
| FT1 | N128 | 65536 | 12 | 12 | 480 | full | 0.62 | 0.93 | 0.26 | 11.20 | 53.25 | 0.47 |
| FT2 | N128 | 65536 | 8 | 8 | 128 | 30 | 0.94 | 0.99 | 0.17 | 0.78 | 5.88 | 0.27 |
| ESM3 | N128 | 4096 | 2 | 30 | 1024 | full | 0.92 | 1.00 | 0.30 | 4.50 | 22.51 | 0.04 |
| FT3 | N128 | 4096 | 8 | 8 | 128 | 30 | 0.92 | 0.99 | 0.26 | 1.16 | 4.42 | 0.41 |
| FT3 | N128 | 1024 | 8 | 8 | 128 | 30 | 0.91 | 0.98 | 0.29 | 1.25 | 4.31 | 0.39 |
| FT3 | N128 | 256 | 8 | 8 | 128 | 30 | 0.89 | 0.98 | 0.22 | 1.49 | 7.76 | 0.37 |
| FT3 | N128 | 128 | 8 | 8 | 128 | 30 | 0.89 | 0.96 | 0.27 | 1.60 | 6.78 | 0.39 |
| FT3 | N128 | 64 | 8 | 8 | 128 | 30 | 0.84 | 0.95 | 0.23 | 2.34 | 15.64 | 0.49 |

##### FoldToken3 works well using 256 or smaller codebook size

FoldToken2 and FoldToken1 utilize a large codebook size of 65,536 to achieve good reconstruction performance. However, we observe that most code vectors are not utilized in the reconstruction process (unbalanced usage), and many code vectors are similar to each other (self-confusion). We attribute the unbalanced usage to the deterministic selection strategy and propose a stochastic sampling selection to give each vector the chance to participate in the learning process. This change increased the codebook usage rate and improved codebook diversity, and also reduced the risk of self-confusion. As a result, FoldToken3 achieves comparable reconstruction performance to FoldToken2 while using a much smaller codebook size of 256, less than 0.4% of FoldToken2. This phenomenon is consistent across single-chain (T116, T493) and multi-chain (M1031) protein reconstruction tasks, even for novel protein structures (N128).

##### When FoldToken3 will Fail on Single-chain Data?

In Fig. 2, we show cases in N128 where the TMScore is less than 0.5. We observe that the model has difficulty reconstructing structures when the protein is too short. This is because proteins with fewer than 30 amino acids were filtered out in the training set. Nevertheless, we can see that the global shape of the protein is still well preserved, although the accuracy of the secondary structure is not satisfactory.

**Figure 2:**
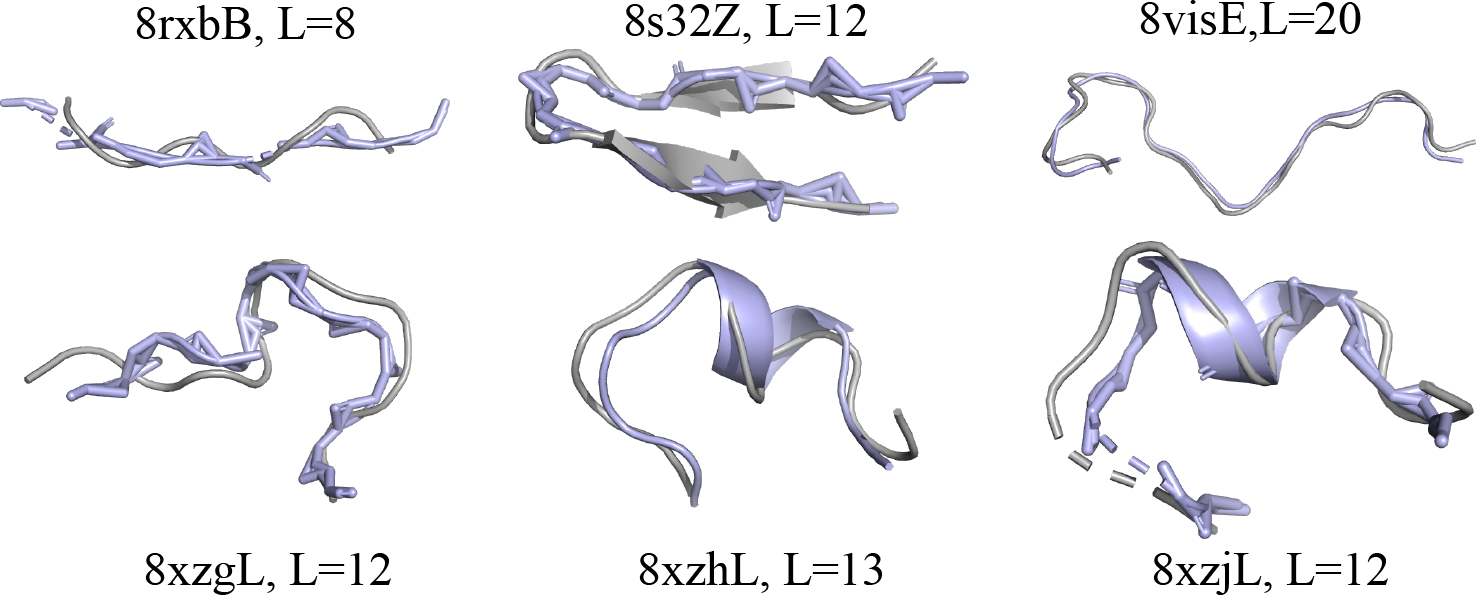
The cases when TMScore<0.5 in N128. Grey structures are the ground truth, and colored structures are the reconstructed ones.

##### FT3 is comparable to ESM3 using less parameters and data

When comparing FoldToken3 with ESM3, we find that FoldToken3 achieves comparable reconstruction performance to ESM3 while using fewer parameters and less data. In terms of trainable parameters, the encoder and decoder of FoldToken3 have **4.31M and 4.92M** parameters, respectively. In comparison, ESM3’s encoder and decoder have **30.1M and 618.6M** parameters, respectively. Regarding the training data, FoldToken3 is trained on the PDB dataset, which is a small subset of ESM3’s training set. Additionally, FoldToken3 is specifically trained for multi-chain protein reconstruction, a more challenging task than the single-chain protein reconstruction that ESM3 is trained for. Nevertheless, the checkpoints learned from the multi-chain task generalize well to single-chain tasks.

#### 3.2 Multi-Chain Benchmark (Q2)

##### Multi-Chain Data for Evaluation

We evaluate the model on the antibody-antigen dataset (SAbDab), which contains 6741 protein complexes. We use foldseek to cluster protein chains into 1323 clusters:

~~~
^1^ foldseek easy - cluster ab_pdb / res tmp -c 0.7
~~~

We use complexes of representative chains from each cluster for evaluation. After filtering representative proteins with less than 30 residues or more than 1000 residues, we get the evaluation dataset containing 1031 protein complexes, named M1031.

In Table. 2, we show the reconstruction results of FoldToken2 and conclude that:

**Table 2:**
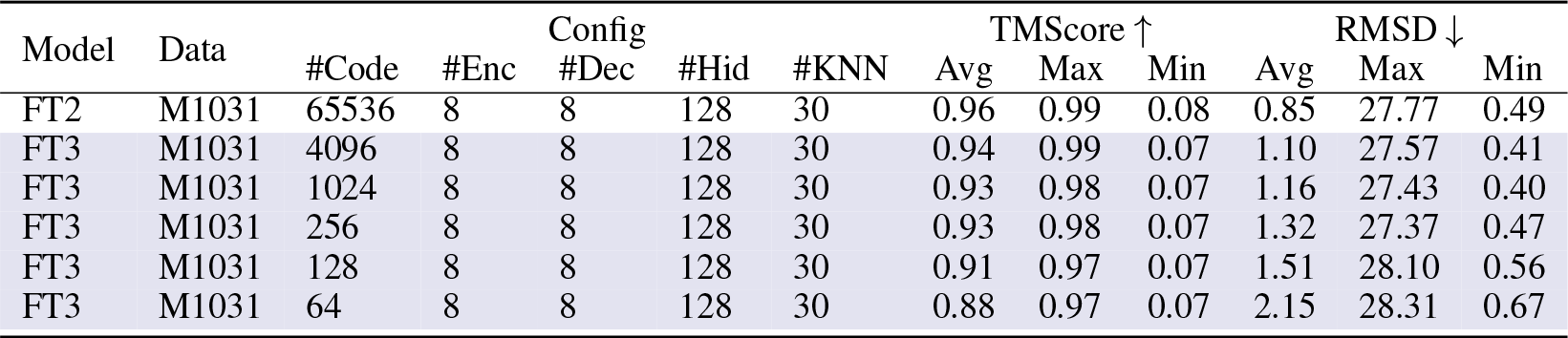
Single-chain Reconstruction Benchmark. FT1 and ESM3 are ignored here, because they cannot handle multi-chain proteins.

##### The compact code space can represent complex interactions

When using 256 code vectors, FoldToken3 can reconstruct protein complex structures effectively, achieving an average TM-score of 0.93 and an average RMSD of 1.32. Previously, we believed that tokenizing single-chain proteins was difficult, especially with very small codebooks. However, FoldToken3 demonstrates that even protein complexes can be represented well using a small codebook. This discovery will promote complex modeling, such as similar interface searching, complex alignment, and complex generation.

##### When FoldToken3 performs worse on multi-chain data?

In Fig. 3, we show the top-9 cases with largest RMSD. The large RMSD is mainly due to the long protein length. The visual examples show that the global&local structures are well preserved in the worst cases. We are suprevised that the model can still reconstruct the protein complex structures effectively, even when the protein is too long and the codebook is too small.

**Figure 3:**
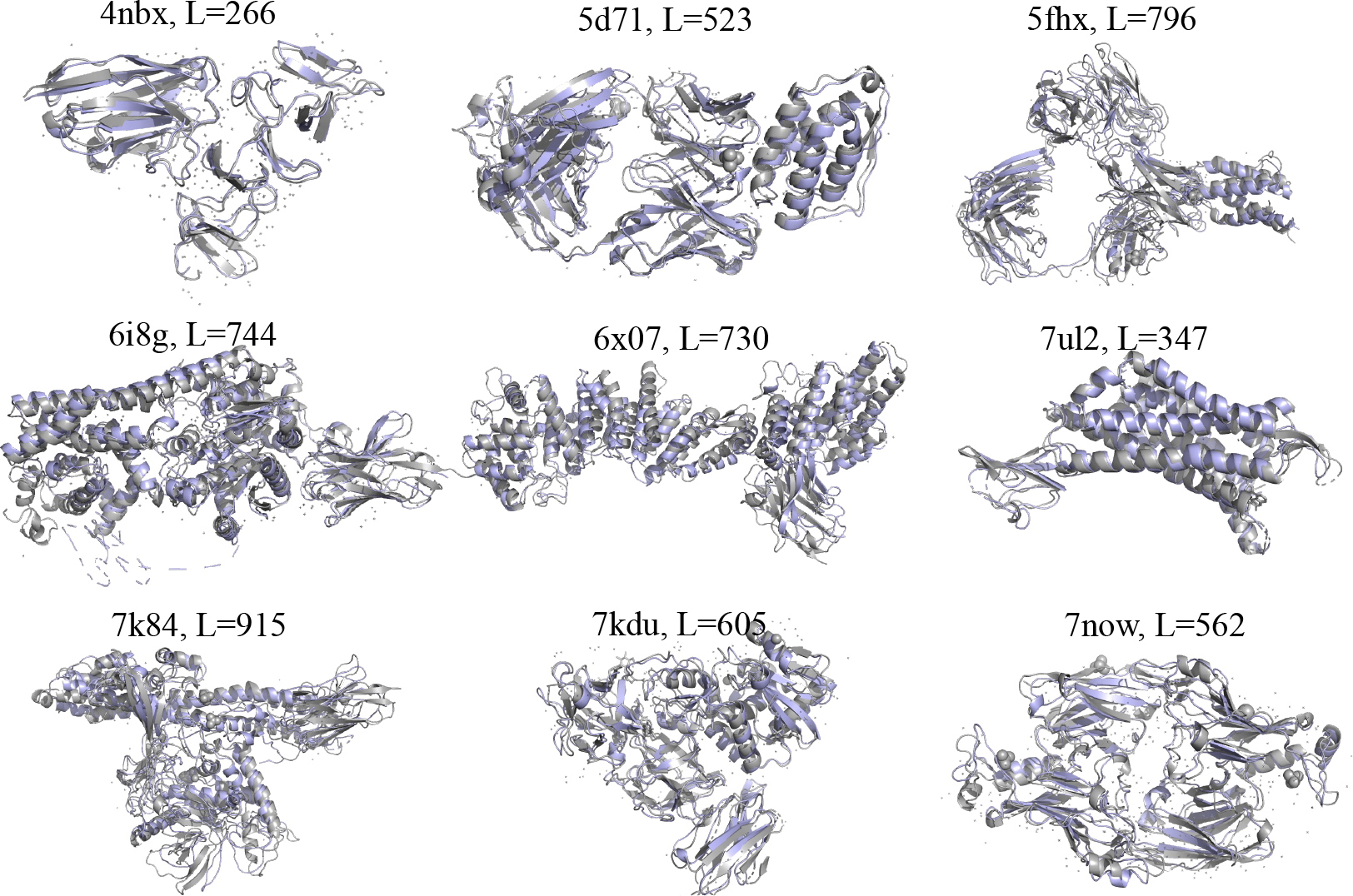
The top-9 cases with largest RMSD in multi-chain setting. Grey structures are the ground truth, and colored structures are the reconstructed ones.

## 4. VQ Insights (Q3)

### Training Stability

In Figure 4, we show the training curves for FoldToken2 and FoldToken3. We observe that when the temperature is very small, the training loss of FoldToken2 crashes due to unstable gradients. However, FoldToken3 introduces the “partial gradient” operation, which helps stabilize the training process. The training crash is also reflected in the similarity metric. For FoldToken2, the similarity approaches 1.0 after the crash, indicating that most code vectors have collapsed to a single vector. In contrast, the similarity of FoldToken3 decreases during training, suggesting that the code vectors remain diverse.

**Figure 4:**
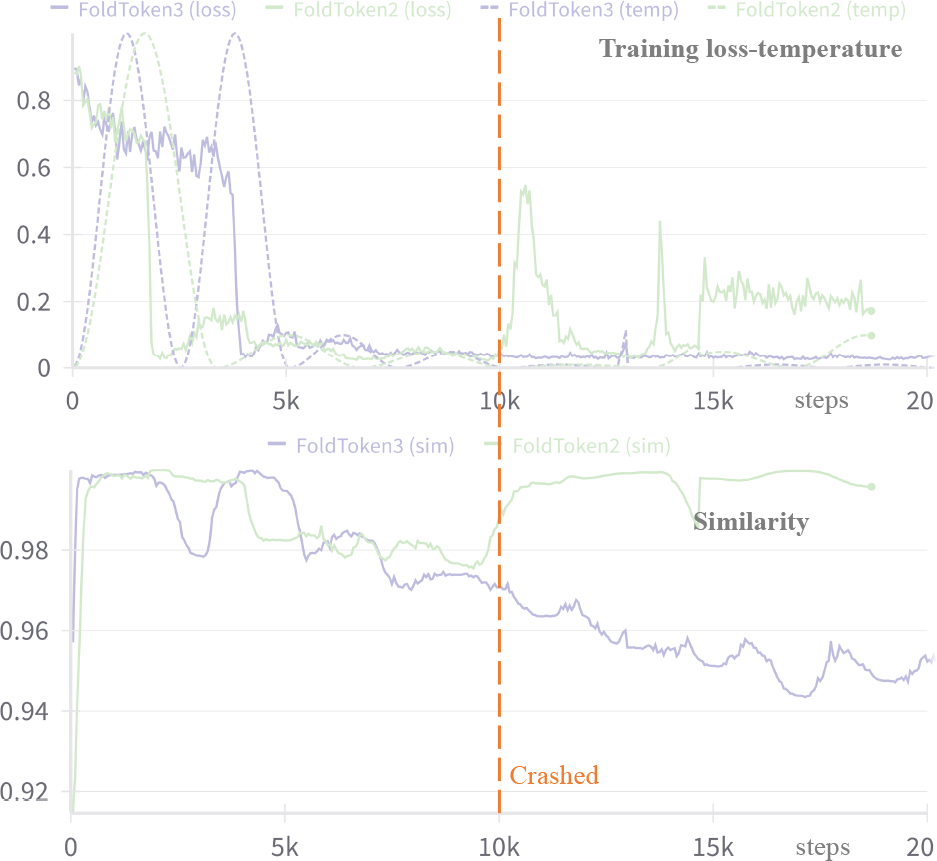
Training curve.

### Code Diversity

We compute the cosine similarity for each code and its nearest neighbor and show the code similarity in Table. 3 and Fig. 5. We observe that the similarity of FoldToken2 is close to 1.0, FoldToken1 has slightly better diversity than FoldToken2, and FoldToken3 has the best diversity. This diversity is crucial for downstream tasks, as it ensures that the model can distinguish different code vectors. The higher the diversity, the lower the risk of confusing two semantic similar tokens.

**Table 3:** The code vector similarity.

| Model | Avg↓ | Max↓ | Min↓ |
| --- | --- | --- | --- |
| FT1 | 0.9959 | 1.0 | 0.7447 |
| FT2 | 0.9999 | 1.0 | 0.9625 |
| FT3 | 0.9070 | 0.9711 | 0.7371 |

**Figure 5:**
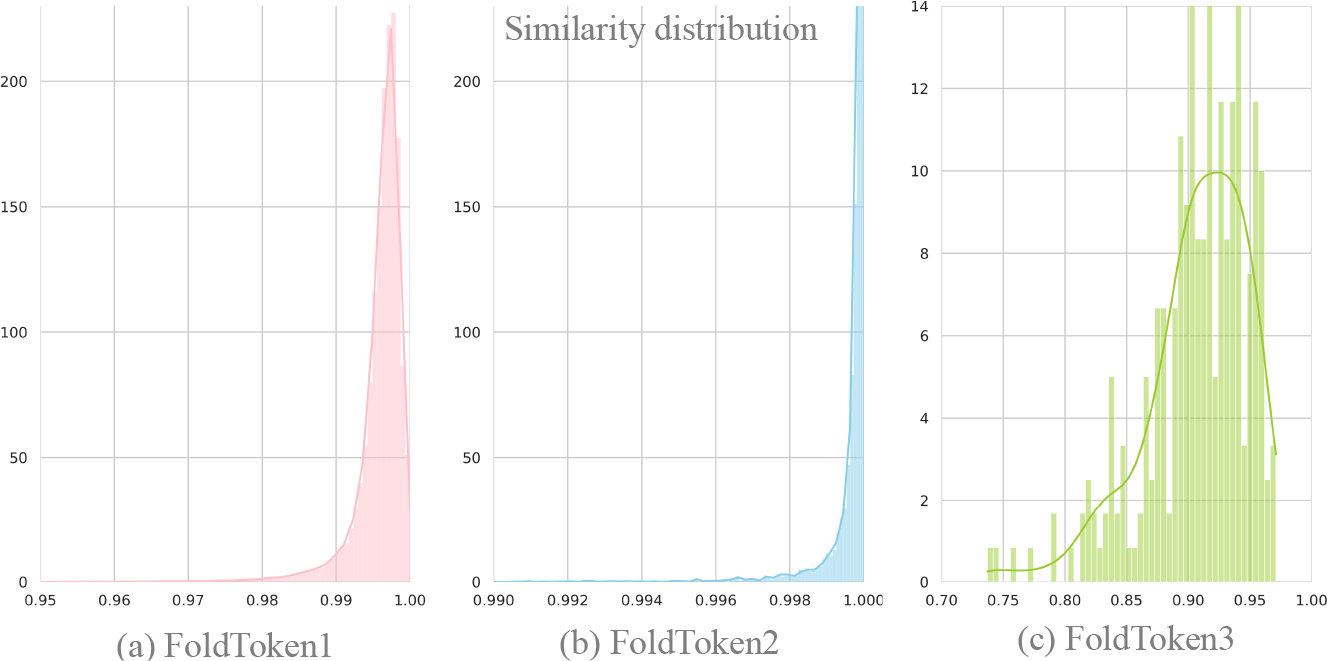
The distribution of code similarity.

### Compression & Utilization Rate

FoldToken2 uses 65,536 code vectors, with a utilization rate of 67.6% over the entire PDB dataset. In comparison, FoldToken3 uses only 256 code vectors, with a 100% utilization rate. This indicates that FoldToken3 can create a more compact discrete representation of protein structures, where the number of used code vectors is only 0.39% of that in FoldToken2. The reduced codebook size will lead to a more consistent and robust fold-language.

## 5 Conclusion

This paper introduces FoldToken3, a novel protein structure tokenization method that can efficiently compress protein structures into 256 or fewer tokens, for both single-chain and multi-chain data, while maintaining reconstruction quality comparable to FoldToken2. To date, FoldToken3 is the most efficient, lightweight, and compression-friendly protein structure tokenization approach. This advancement will benefit a wide range of protein structure-related tasks, including protein structure alignment, generation, and representation learning.

